# *In vivo* nuclear RNA structurome reveals RNA-structure regulation of mRNA processing in plants

**DOI:** 10.1101/839506

**Authors:** Zhenshan Liu, Qi Liu, Xiaofei Yang, Yueying Zhang, Matthew Norris, Xiaoxi Chen, Jitender Cheema, Yiliang Ding

## Abstract

mRNA processing is critical for gene expression. A challenge in regulating mRNA processing is how to recognize the actual mRNA processing sites, such as splice and polyadenylation sites, when the sequence content is insufficient for this purpose. Previous studies suggested that RNA structure affects mRNA processing. However, the regulatory role of RNA structure in mRNA processing remains unclear. Here, we performed *in vivo* selective 2’-hydroxyl acylation analysed by primer extension (SHAPE) chemical profiling on *Arabidopsis* nuclear RNAs and generated the *in vivo* nuclear RNA structure landscape. We found that nuclear mRNAs fold differently from cytosolic mRNAs. Notably, we discovered a two-nucleotide single-stranded RNA structure feature upstream of 5’ splice sites that is strongly associated with splicing and the selection of alternative 5’ splice sites. Moreover, we found the single-strandedness of branch point is also associated with 3’ splice site recognition. We also identified an RNA structure feature comprising two close-by single-stranded regions that is specifically associated with both polyadenylation and alternative polyadenylation events. Our work demonstrates an RNA structure regulatory mechanism for mRNA processing.

## Introduction

In eukaryotes, mRNAs undergo several processing steps including 5′ capping, splicing, and 3′ cleavage/polyadenylation to become functional mature mRNAs. Thus, mRNA processing plays a critical role during gene expression^1,2^. Over past decades, a key question is how mRNA processing sites, such as polyadenylation and splice sites, are precisely recognized in the transcriptome, particularly from surrounding sites with similar sequence content^3,4^. For instance, 5’ splice site recognition was found to be not always dependent on the sequence content of U1 snRNA binding motif. Some 5’ splice sites were selected over those flanking sites with better complementarity to U1 snRNA binding sequence^4^. In case-by-case studies, quite a number of RNA binding proteins have been identified that contribute to the recognition of actual polyadenylation and splice sites^4,5^. However, a general regulatory mechanism that recognizes actual sites during mRNA processing is lacking. As an intrinsic characteristic of RNA molecules, RNA structure was suggested to be involved in mRNA processing^6^. Previous individual studies suggested that RNA structure can affect polyadenylation and splicing^7–13^. Yet, how RNA structure contributes to the recognition of polyadenylation and splice sites, in general, remains elusive.

With recent advances in RNA structure profiling^14–16^, more attention has been drawn toward understanding how RNA structure influences mRNA processing. Previous *in vitro* enzymatic RNA structure profiling (utilizing RNases that selectively cleave either single-stranded or double-stranded nucleotides) in *Arabidopsis* nuclear RNAs, found that the 5’ end of introns were more double-stranded compared to upstream exons, and the 3’end of introns were more single-stranded compared to upstream intron regions^14^. However, no significant structure signatures were identified for either polyadenylation or alternative polyadenylation sites^14^. This may be due to limitations imposed by using RNases, which are quite bulky and less sensitive in detecting specific RNA structures, compared to the relatively small chemicals used for RNA structure probing^17,18^. Furthermore, several previous studies have shown that *in vitro* RNA structures were not able to reflect the proper folding status of RNAs in living cells^19,20^. A recent *in vivo* dimethyl sulfate (DMS) RNA structure profiling study on human mature mRNAs identified RNA structure features for polyadenylation (poly(A)) sites^15^. A more folded structure downstream of the polyadenylation signal motif was identified that facilitated polyadenylation^15^. However, mammalian RNAs were found to adopt different structure conformations in different cellular compartments^21^. Thus, the structure of mature mRNAs in the cytosol is likely to be different from the structure of pre-mRNA in the nucleus. If so, mature mRNA structures are unlikely to reveal the role of RNA structure in polyadenylation. A notable limitation of this DMS method is the loss of RNA structure information for the half transcriptome because DMS only detects structure information of As and Cs, lacking the base-pairing status of Us and Gs.

Here, we studied the role of RNA structure in mRNA processing by performing *in vivo* SHAPE (**S**elective 2’ **H**ydroxyl **A**cylation analysed by **P**rimer **E**xtension) chemical probing on *Arabidopsis thaliana* nuclear RNAs, to generate the first *in vivo* RNA structure landscape with all four nucleotides in plants. We found that nuclear mRNA structures are globally different from cytosolic mRNA structures in *Arabidopsis*. Our study further successfully dissected pre-mRNA structure features before mRNA processing and determined the regulatory role of RNA structure during mRNA maturation.

## Results

### Nuclear SHAPE-Structure-Seq generates *in vivo* RNA structure landscape of *Arabidopsis* nuclear RNAs with high coverage and accuracy

To investigate the role of RNA structure in mRNA processing, we performed SHAPE chemical probing^22^ on *Arabidopsis* nuclear RNAs and generated the first *in vivo* RNA structure profiles with all four nucleotides in plants. Firstly, SHAPE reagent (2-methylnicotinic acid imidazolide, NAI) treatment was applied on 5-day-old *Arabidopsis* seedlings^22^ (Fig. 1a). Intact nuclei were isolated and nuclear RNAs were extracted. The intactness of isolated nuclei was confirmed by microscopy imaging with DAPI staining^23^ (Supplementary Fig. 1a). Enrichment of nuclear histone H3 protein and absence of cytoplasmic protein PEPC (Phosphoenolpyruvate carboxylase) in the isolated nucleus, further confirmed the high purity and quality of the isolated nuclei (Supplementary Fig. 1b). We generated two independent biological replicates of (+)SHAPE (samples with SHAPE treatment) and (-)SHAPE (control samples without SHAPE treatment) Structure-Seq libraries for high-throughput sequencing^24,25^ (Fig. 1a and Supplementary Fig. 2, see Methods). Given that interactions between RNA and RNA binding proteins can prevent the SHAPE modification^22^, we also performed SHAPE treatment on nuclear RNAs after removing proteins thus generating *deproteinized* nuclear SHAPE-Structure-Seq libraries in parallel (Fig. 1a) to assess any effect on SHAPE modification signals caused by protein protection. The *deproteinized* libraries were designed to preserve RNA secondary structure after cell lysis and protein removal but not subjected to RNA denaturing under high temperature. Thus, the *deproteinized* condition here is still *in vitro* condition. Over 616 million 100bp paired-end reads per library were generated and further mapped onto *Arabidopsis* genome sequences (TAIR10) with additional alternative spliced isoforms annotated from AtRTD2 database^26^ (Supplementary Table 1).

**Fig. 1.**
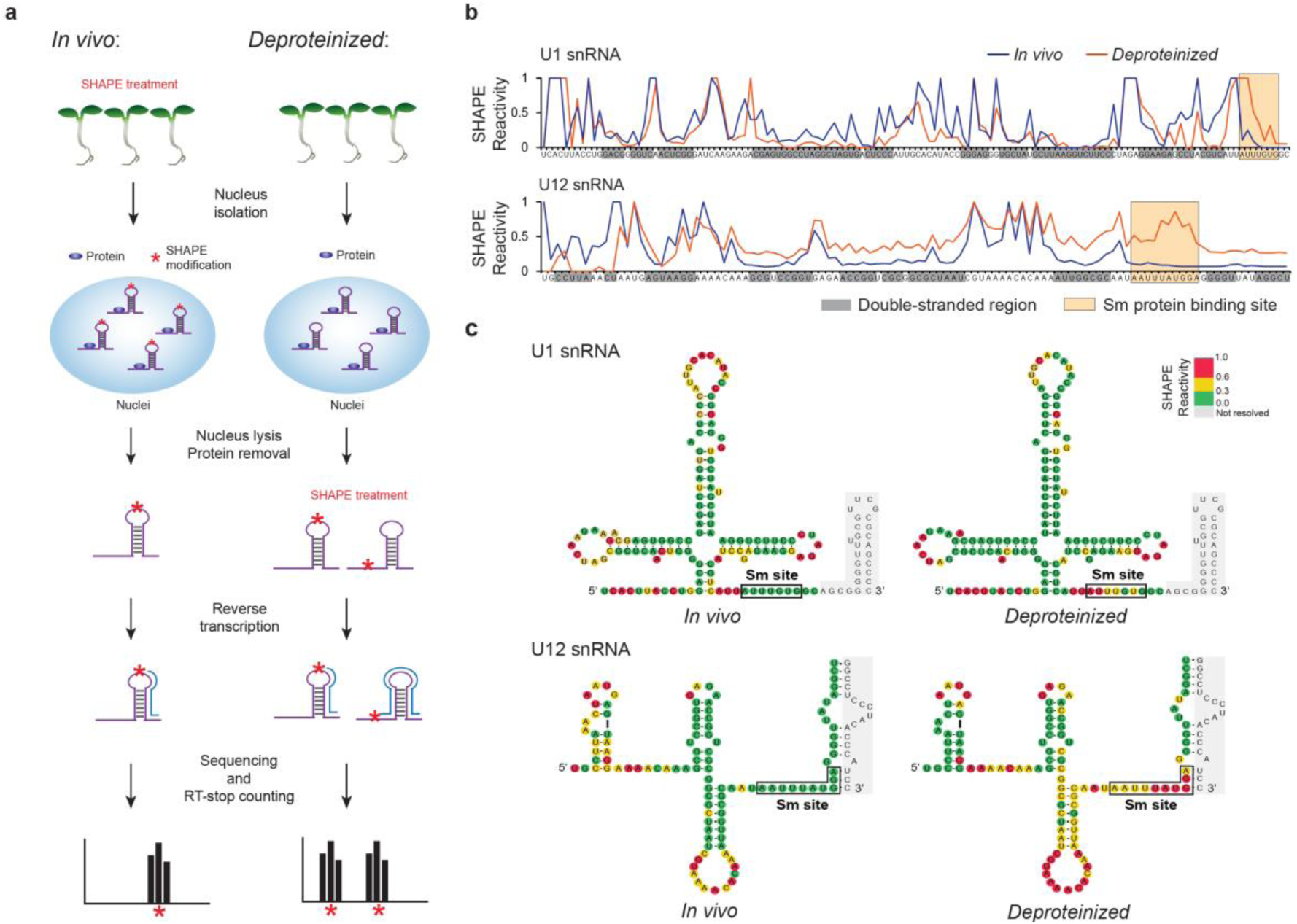
SHAPE-Structure-Seq method can accurately probe the *in vivo* RNA structure of nuclear RNAs. **a**, Schematic pipeline of nuclear SHAPE-Structure-Seq for both *in vivo* and *deproteinized* conditions. Asterisks, SHAPE modification; blue oval, protein; RT, reverse transcription. For *in vivo* treatments (left), NAI was applied to *Arabidopsis thaliana* seedlings directly and single-stranded nucleotides of RNA were modified. SHAPE treatment was also applied on the RNAs after removing protein, which we termed the ‘*deproteinized* condition’ (right). Deep sequencing was performed followed by the RT-stop counting. **b**, SHAPE reactivity profiles of U1 and U12 snRNAs. SHAPE reactivity profiles of both *in vivo* (blue) and *deproteinized* (orange) conditions were shown. Double-stranded regions were shaded with grey. Sm protein binding sites were highlighted with yellow boxes. At Sm protein binding sites, significantly higher SHAPE reactivities were observed under the *deproteinized* condition rather than the *in vivo* condition for both U1 and U12 snRNAs (Paired *t*-test, *P-value*= 6.8e-3 and 3.1e-6 for U1 and U12 snRNA respectively). Higher *deproteinized* SHAPE reactivities were also observed at some double-stranded regions of U12 snRNA, suggesting the structure of these regions might also be affected by protein interaction. **c**, SHAPE reactivities are consistent with the phylogenetically derived U1 and U12 snRNA structures. Sm protein binding sites were highlighted with black boxes. Nucleotides were colour-coded according to *in vivo* and *deproteinized* SHAPE reactivity values (SHAPE reactivity 0.6-1.0 marked in red, 0.3-0.6 marked in yellow, 0-0.3 marked in green).

Nucleotide modification in both (+)SHAPE and (-)SHAPE libraries were highly concordant, with slight enrichment in (+)SHAPE shown for As and Us over Cs and Gs, as expected, since As and Us tend to be more single-stranded than Cs and Gs (Supplementary Fig. 3a). The high correlation of mRNA abundance between the two biological replicates indicated the high reproducibility of our nuclear SHAPE-Structure-Seq libraries (Supplementary Fig. 3b). To further validate the reproducibility of our SHAPE structure probing, we compared SHAPE reactivity profiles of U1 and U12 snRNA between the two biological replicates and noted a high correlation between them (Pearson correlation coefficient=0.93-0.97) (Supplementary Fig. 3c). Thus, we merged these two biological replicates for further RNA structure analysis.

We assessed both the sequencing reads coverage and reverse-transcription stop counts of our nuclear SHAPE-Structure-Seq libraries. Notably, more than 20,752 genes had at least 10 reads per nucleotide coverage (Supplementary Fig. 4a), among which more than 12,366 genes reached the threshold of at least one reverse-transcription stop count per nucleotide for RNA structure analysis (Supplementary Fig. 4b). To assess the accuracy of our RNA structure profiling, we compared SHAPE reactivity profiles of U1 and U12 snRNAs with their phylogenetically derived structures, which are evolutionarily conserved structures and are the closest models of *in vivo* structure^22,27^. Overall, the SHAPE reactivities were consistent with phylogenetically derived RNA structures where high SHAPE reactivities were observed in single-stranded regions, while low SHAPE reactivities were at double-stranded nucleotides (Fig. 1b,c, Supplementary Table 2). Both U1 and U12 snRNAs interact with Sm proteins to form small nuclear ribonucleoparticle structures^22,27^. We also found that SHAPE reactivities at Sm protein binding sites of U1 and U12 snRNA were significantly higher in the *deproteinized* rather than *in vivo* condition (Fig. 1b,c). In addition, global SHAPE reactivities were also found to be significantly higher in the *deproteinized* condition compared to the *in vivo* condition suggesting that absence of protein protection in the *deproteinized* condition allowed nucleotide modification by SHAPE (Supplementary Fig. 5). Collectively, these results indicated that our nuclear SHAPE-Structure-Seq method can accurately probe *in vivo* RNA structures of nuclear RNAs.

### Nuclear mRNA structures are globally different from cytosolic mRNA structures

Cytosolic mRNAs are the processed products from nuclear mRNAs, thus they share the same sequences. However, whether they share the same RNA structure features remains unclear. To address this question, we generated *in vivo* SHAPE-Structure-Seq libraries of *Arabidopsis* cytosolic mRNAs in parallel. We then compared these libraries with our *in vivo* nuclear SHAPE-Structure-Seq libraries (Supplementary Data 1 and 2). Firstly, we compared average SHAPE reactivities of exons between nuclear and cytosolic mRNAs. We found that exons in cytosolic mRNAs had significantly higher average SHAPE reactivities than those in nuclear mRNAs, suggesting exons in cytosolic mRNAs tended to be more single-stranded than those in nuclear mRNAs (Fig. 2a). This result is also similar with that observed in mammals^21^. We further compared average SHAPE reactivities in different genic regions of exons: the 5’ untranslated region (5’UTR), the coding region (CDS) and the 3’ untranslated region (3’UTR), between nuclear and cytosolic mRNAs. Notably, we found that average SHAPE reactivities in both 5’UTR and 3’UTR were significantly higher in nuclear mRNAs than those in cytosolic mRNAs (Fig. 2b). In contrast, significantly lower average SHAPE reactivities were observed in nuclear mRNA CDS regions compared to those in cytosolic mRNAs (Fig. 2b). Previous studies on total mRNAs dominated by cytosolic mRNAs observed unique structure features across translation start and stop codons that were associated with translation^24,28–30^. Consistent with these observations, we also found higher SHAPE reactivities upstream of start codons, lower SHAPE reactivities downstream of start codons, and higher SHAPE reactivities at stop codons compared to flanking regions (Fig. 2c,d) in our cytosolic SHAPE-Structure-Seq libraries. We then compared average SHAPE reactivity profiles between nuclear and cytosolic mRNAs across these two sites. Significantly higher SHAPE reactivities downstream of start codons and significantly lower SHAPE reactivities at stop codons in nuclear mRNAs were observed, compared to those in cytosolic mRNAs (Fig. 2c,d). Taken together, nuclear mRNA structures are globally different from cytosolic mRNA structures, which implies nuclear and cytosolic mRNAs might adopt different structures to serve their respective biological functions, e.g. translation in the cytosol and mRNA processing in the nucleus. Therefore, we further investigated how nuclear mRNA structures are associated with mRNA processing.

**Fig. 2.**
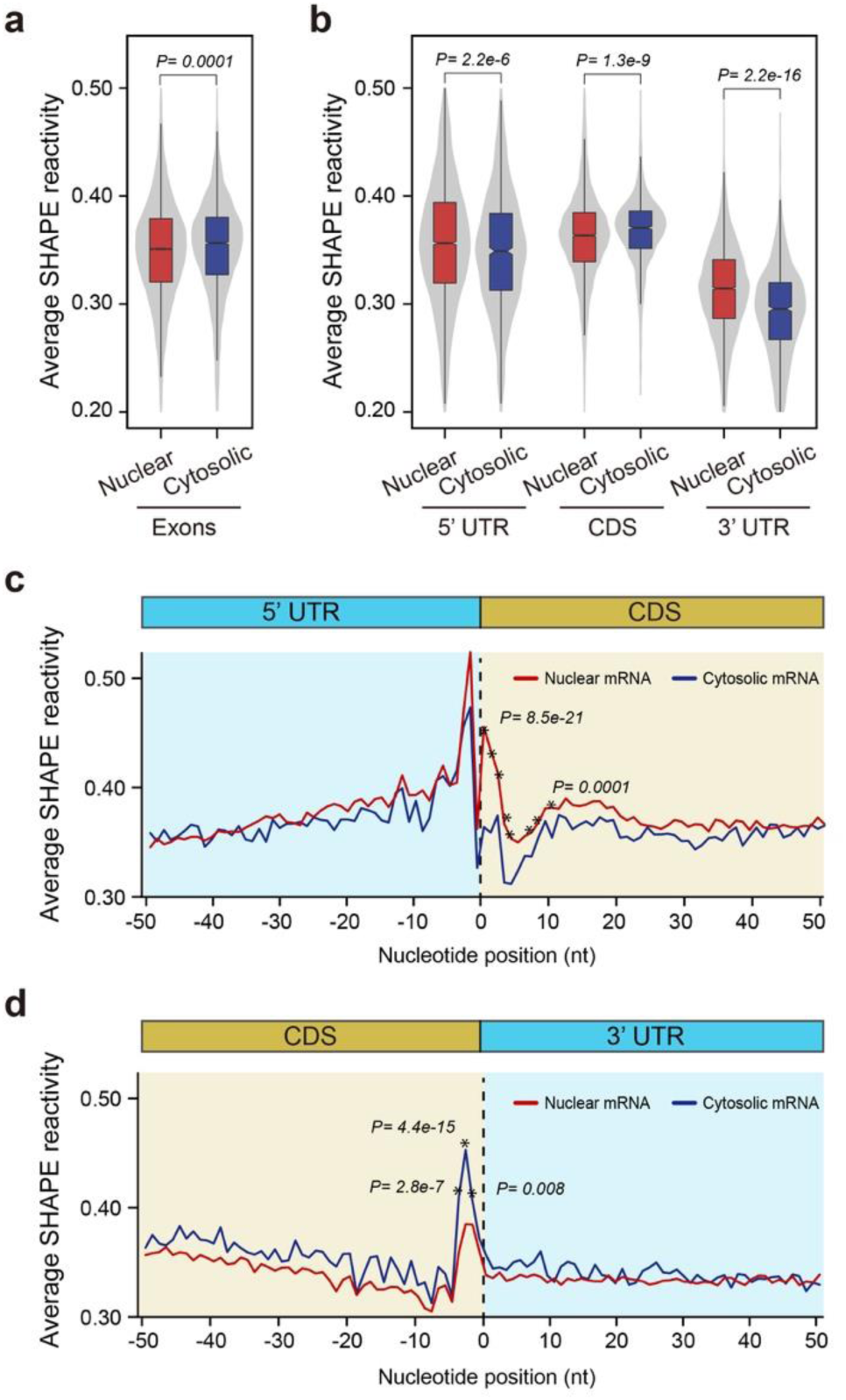
*In vivo* nuclear mRNA structures are globally different from cytosolic mRNA structures. **a**, Comparison of SHAPE reactivities between the exon regions of nuclear and cytosolic mRNAs. The average SHAPE reactivity of exons in nuclear mRNAs is significantly lower than that in cytosolic mRNAs (Mann-Whitney test, the *P-value* is shown). **b**, Comparisons of average SHAPE reactivities between nuclear and cytosolic mRNAs for 5’ UTR, CDS and 3’UTR. Average SHAPE reactivities in both 5’UTR and 3’UTR are significantly higher in nuclear mRNAs than those in cytosolic mRNAs, whereas average SHAPE reactivities in CDS are significantly lower in nuclear mRNA than those in cytosolic mRNAs (Mann-Whitney test, the *P-values* were shown). **c**, Comparison of average SHAPE reactivity profiles between nuclear and cytosolic mRNAs across the translation start codon. Average SHAPE reactivities downstream of the start codon are significantly higher in nuclear mRNAs compared to cytosolic mRNAs (Mann-Whitney test, the highest and lowest *P-*values for the first ten nucleotides of the CDS region are shown). **d**, Comparison of SHAPE reactivity profiles between nuclear and cytosolic mRNAs across the translation stop codon. Average SHAPE reactivities at the stop codon are significantly lower in nuclear mRNAs compared to cytosolic mRNAs (Mann-Whitney test, the *P-values* were shown).

### Distinctive pre-mRNA structure features are strongly associated with both splicing and alternative splicing

Splicing is a key mRNA processing step that was previously suggested to be influenced by RNA structure^8^. Since only pre-mRNA structure before splicing (unspliced primary transcripts) can be used for dissecting the mechanism underpinning splicing, we firstly assessed whether pre-mRNAs were enriched in our nuclear SHAPE-Structure-Seq data. We found that the expression abundance of constitutively spliced introns was much higher in our nuclear SHAPE-Structure-Seq libraries compared to our cytosolic SHAPE-Structure-Seq libraries, indicating high enrichment of pre-mRNAs in our nuclear SHAPE-Structure-Seq data (Supplementary Fig. 6). Since nuclear mRNAs still contain spliced transcripts, we only used reads mapped across exon-intron junctions and in intron regions for generating SHAPE reactivity profiles to obtain accurate RNA structure information of pre-mRNAs before splicing (See details in Methods). Also, to eliminate any ambiguous reads assignment at the conserved dinucleotide AG at 3’ splice sites (3’ss), we only calculated SHAPE reactivities across 5’ splice site (5’ss) and the whole intron except for AG at 3’ss (See details in Methods).

In addition to generating RNA structure information of pre-mRNAs, we also calculated the splicing efficiency for each intron to measure the outcome for splicing events (Supplementary Fig. 7, See details in Methods). Since most of the introns showed either very high (>=90%) or very low (<=10%) splicing efficiencies, two groups of splicing events were classified: spliced events (splicing efficiency >= 90%, 32,522 spliced events were identified, Supplementary Data 3) and unspliced events (i.e. intron retention, splicing efficiency <= 10%, 4,056 unspliced events were identified, Supplementary Data 3). We compared average SHAPE reactivity profiles between these two groups of splicing events. Although the exon-intron regions of both spliced and unspliced events shared similar nucleotide compositions (Supplementary Fig. 8), distinctive SHAPE reactivity profiles were observed between these two groups (Fig. 3a,b). Specifically, we found that *in vivo* SHAPE reactivities at the −1 position immediately upstream of 5’ss were notably higher for spliced events compared to unspliced events (Fig. 3a). Similarly, SHAPE reactivities at the −1 and −2 positions upstream of 5’ss were significantly higher in spliced events than those in unspliced events for the *deproteinized* condition. These findings indicated that the −1 and −2 nucleotides upstream of 5’ss tended to be more single-stranded in spliced events compared to unspliced events (Fig. 3b). We further assessed sequence content across 5’ss in both spliced and unspliced events and found no sequence preference between these two groups (Supplementary Fig. 8). Thus, our results suggested that this distinctive structure signature was associated with splicing events, but not due to sequence preference.

**Fig. 3.**
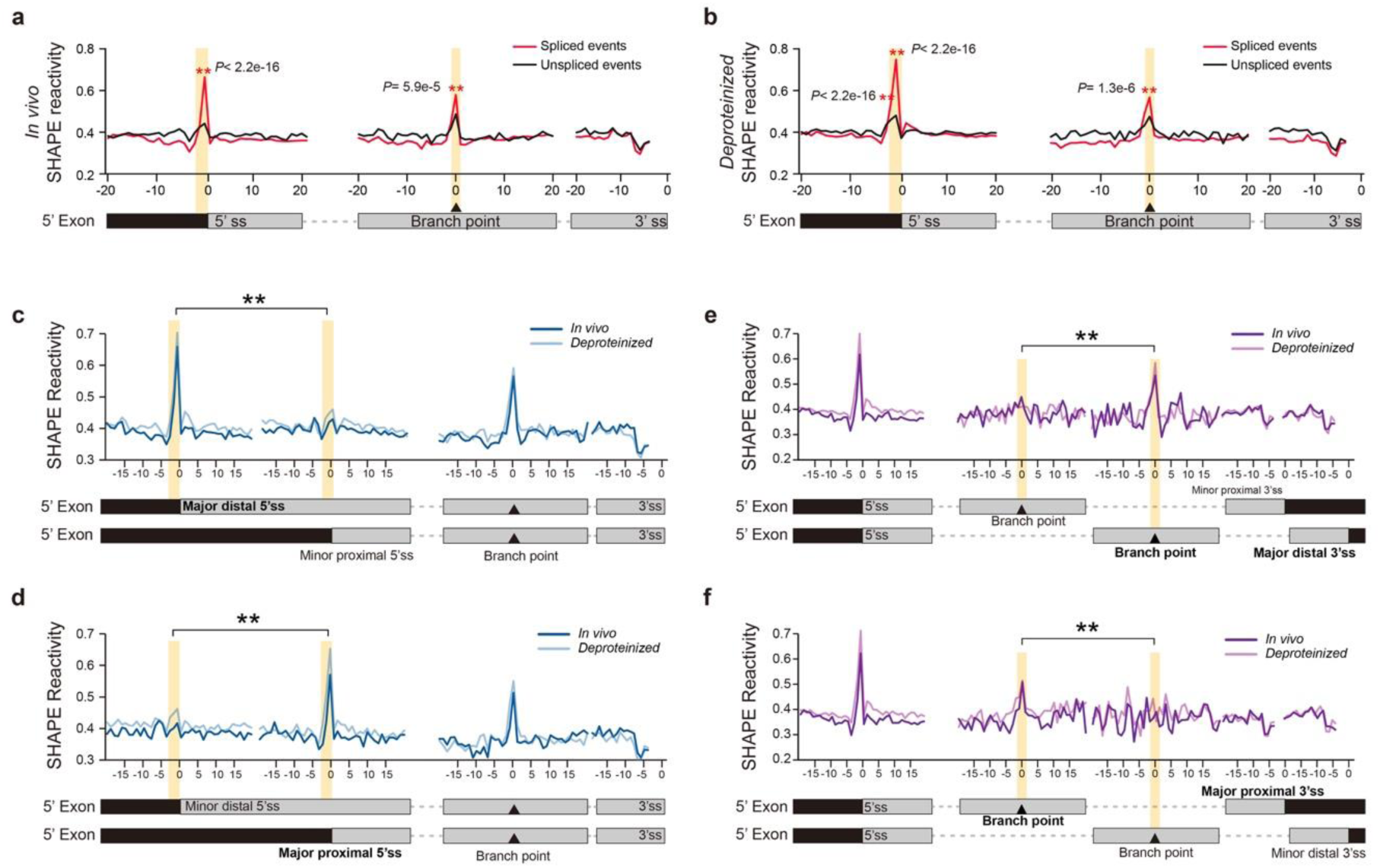
pre-mRNA secondary structure features upstream of 5’ss and at the branch site are associated with splicing and alternative splice site selection. **a, b**, SHAPE reactivity profiles across 5’ss, branch point and 3’ss for *in vivo* (**a**) and *deproteinized* (**b**) conditions. Average SHAPE reactivity profiles for spliced (red) versus unspliced (black) events are shown. Significantly higher SHAPE reactivities are observed at the −1 and −2 nt positions of 5’ss and the branch point for spliced events rather than unspliced events (Marked with asterisks, Mann-Whitney test, *P*-values are shown). **c**, SHAPE reactivity profiles for alternative 5’ss events with the distal 5’ss as the major one. Average SHAPE reactivity profiles of both *in vivo* (dark blue) and *deproteinized* (light blue) conditions are shown. Gene models for the two alternative isoforms are shown at the bottom. Significantly higher SHAPE reactivities only appear at −1 and −2 positions upstream of the major distal 5’ss rather than the minor proximal 5’ss (Mann-Whitney test, *P*-value = 1.6e-4 and <2.2e-16 at −1 and −2 positions under *in vivo* condition; *P*-value = 6.1e-9 and <2.2e-16 at −1 and −2 positions under *deproteinized* condition). **d**, SHAPE reactivity profiles for alternative 5’ss events with the proximal 5’ss as the major one. The significantly higher SHAPE reactivities of −1 and −2 positions only appear upstream of the major proximal 5’ss rather than the minor distal 5’ss (Mann-Whitney test, *P*-value = 3.3e-12 at −1 position under *in vivo* condition; no significant difference was detected at −2 position under *in vivo* condition; *P*-value = 3.1e-5 and <2.2e-16 at −1 and −2 positions under *deproteinized* condition). **e**, SHAPE reactivity profiles for alternative 3’ss events with the distal 3’ss as the major one. Average SHAPE reactivity profiles of both *in vivo* (dark purple) and *deproteinized* (light purple) conditions across different 3’ss and the corresponding branch points are shown. Significantly higher SHAPE reactivity only appears at the branch point of the major distal 3’ss rather than the minor proximal 3’ss (Mann-Whitney test, *P*-value = 1.2e-3 and 2.8e-4 at branch point under *in vivo* and *deproteinized* conditions respectively). **f**, SHAPE reactivity profiles for alternative 3’ss events with the proximal 3’ss as the major one. The significantly higher SHAPE reactivity only appears at the branch point of the major proximal 3’ss rather than the minor distal 3’ss (Mann-Whitney test, *P*-value = 1.4e-2 and 1.7e-3 at the branch point under *in vivo* and *deproteinized* conditions respectively).

We then assessed RNA structure features for branch sites and 3’ss regions, which are important for 3’ss recognition during splicing^1^. To assess RNA structure features at branch sites, we predicted branch sites using SVM-BPfinder^31^. Higher SHAPE reactivities were observed at branch points under both *in vivo* and *deproteinized* conditions for spliced events compared to unspliced events, indicating single-strandedness at the branch point was associated with splicing (Fig. 3a,b). SHAPE reactivities of regions immediately upstream of dinucleotide AG at 3’ss (from −7 to −4 positions) were relatively lower than flanking regions (Fig. 3a,b). However, there was no significant SHAPE reactivity difference between spliced and unspliced events at 3’ss regions, indicating no direct association with splicing. Therefore, both RNA structure features upstream of 5’ss and at the branch point in pre-mRNAs were associated with splicing.

We then explored whether these RNA structure features are also associated with splice site selection in alternative splicing events. Firstly, we identified alternative 5’ss events from genome annotation and selected those pre-mRNAs with two alternative 5’ss (5,116 alternative 5’ss events were identified and used in the following analysis, Supplementary Data 4). We then classified these two alternatives 5’ss as being either distal or proximal 5’ss, according to their relative positions. Based on the expression levels of the corresponding isoforms, we then identified major 5’ss (>=80% of the total abundance of two isoforms) and minor 5’ss (<=20% of the total abundance of two isoforms) (See details in Method). We found that SHAPE reactivities at the −1 and −2 positions upstream of 5’ss were significantly higher in the major 5’ss group than those in the minor 5’ss group, regardless of distal or proximal positions (Fig. 3c,d). Therefore, the two-nucleotide single-stranded RNA structure feature upstream of 5’ss was associated with the selection of alternative 5’ss. We then performed the corresponding assessment for alternative 3’ss events (9,237 alternative 3’ss events were identified and used in the following analysis, Supplementary Data 5) and found SHAPE reactivity at the branch point was notably higher in the major 3’ss group compared to the minor 3’ss group, regardless of distal or proximal positions (Fig. 3e,f). Thus, single-strandedness at the branch point was associated with the selection of alternative 3’ss. High SHAPE reactivity peaks were also observed at other positions around the branch point in both major and minor 3’ss groups, suggesting these high SHAPE reactivities did not contribute to 3’ss selection (Fig. 3f). Taken together, RNA structure features identified upstream of 5’ss and at the branch point were also strongly associated with the recognition of alternative 5’ss and 3’ss in alternative splicing events.

### The two-nucleotide single-stranded RNA structure feature upstream of 5’ss is sufficient to regulate splicing

A nucleotide with high GC content tends to be more double-stranded^32^. Thus, the distinctive single-strandedness at the −1 nucleotide upstream of 5’ss, as a conserved G, is unexpected. In addition, the −1 and −2 nucleotide positions lie within the nine nucleotide binding region of U1 snRNA (from −3 to +6 nt region of 5’ss) during splicing^33^. If this splicing associated RNA structure feature we observed, affected U1 snRNA binding, then a similar RNA structure feature should have been observed across the whole binding site. However, high SHAPE reactivities were only observed for two out of nine nucleotides rather than the whole binding site. Consequently, we tested whether these two single-stranded nucleotides upstream of 5’ss were sufficient to regulate splicing. We selected the first exon-intron-exon region of *AT5G56870* successfully spliced as a representative example of the pre-mRNAs comprising this distinctive two-single-stranded RNA structure feature upstream of 5’ss (Fig. 4a). We then made use of it for our functional validation. To avoid disrupting base-pairing between 5’ss and U1 snRNA during splicing, we maintained the U1 snRNA binding site sequence content and inserted a short sequence immediately upstream of this U1 binding site to form a stable hairpin structure with the whole U1 binding site completely base-paired (illustrated in Fig. 4b). Then, we introduced a series of mutations in the inserted sequence that base-pair with the U1 binding site in order to disrupt the base-pairing status of different nucleotides within this binding region (Fig. 4b). We assessed the splicing events on these designed constructs through transient expression assays in *Nicotiana benthamiana* (Fig. 4c). First, we confirmed that the native sequence construct was successfully spliced in tobacco leaves (Fig. 4c, lane 1). Splicing was completely inhibited when the whole U1 snRNA binding site was completely base-paired with the inserted sequence upstream (Fig. 4c, lane 2). By introducing a mutation “AA” to allow base-pairing disruption at −1 and −2 positions upstream of 5’ss, we found splicing was rescued (Fig. 4c, lane 3). To avoid potential effects due to changing the sequence content, we also mutated these two nucleotides to “GG” that also disrupted the base-pairing status at −1 and −2 positions and found splicing was also rescued (Fig. 4c, lane 4). Furthermore, we assessed the other mutations designed to disrupt other base-pairing sites across the whole U1 binding site (Fig. 4c, lane 5-12). Remarkably, structure disruptions of all other base-pairing sites, even a three-nucleotide mutation, were not able to rescue splicing (Fig. 4c, lane 5-12). Hence, our results indicated that only the two-nucleotide single-stranded RNA structure feature at −1 and −2 positions upstream of 5’ss was sufficient to regulate splicing.

**Fig. 4.**
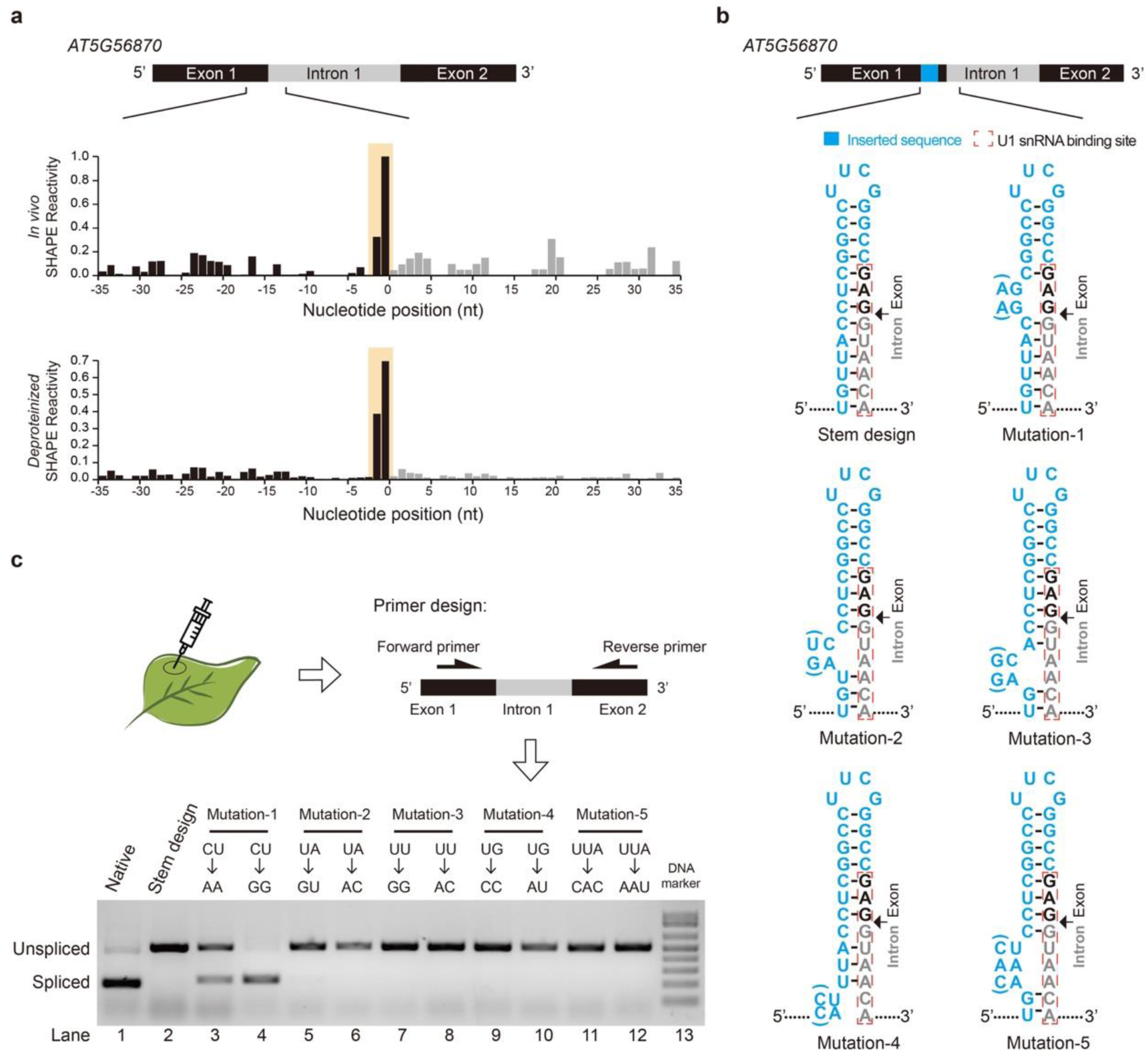
The two-nucleotide single-stranded RNA structure feature at −1 and −2 nt positions upstream of 5’ss can regulate splicing. **a**, SHAPE reactivity profiles across 5’ss of the first intron of *AT5G56870*. High SHAPE reactivities are observed at −1 and −2 nt positions (shaded in yellow) upstream of 5’ss under both *in vivo* (top) and *deproteinized* (bottom) conditions, which resemble the global SHAPE reactivity profiles for spliced events. **b**, Schematic of experimental design to validate the effect of single-strandedness at the −1 and −2 positions of 5’ss on splicing. A short sequence (blue) was inserted immediately upstream of the U1 snRNA binding site (red dashed box) to form a stable hairpin structure with the whole U1 binding site completely base-paired. The exon and intron sequences are colored in black and grey respectively. A series of mutations were introduced at different positions of the inserted sequence to disrupt the base-pairing status of different nucleotides within the U1 binding site. Two types of mutations (with/without bracket) were designed for each position to avoid potential effects due to changing the sequence content. **c**, Determination of splicing events by transient expression assay in *Nicotiana benthamiana*. The spliced and unspliced products were distinguished by semi-qPCR using the same pair of primers located upstream and downstream of the intron. Spliced and unspliced products are indicated by bands with different sizes. The construct with native sequence was successfully spliced (lane 1). The splicing was completely inhibited in the stem design (lane 2). The mutation “AA” or “GG” disrupted the base-pairing status at −1 and −2 positions upstream of 5’ss (Mutation-1) and rescued the splicing (lane 3 and 4). All other mutations (Mutation 2-5) designed to disrupt other base-pairing sites across the U1 binding site did not rescue the splicing (lanes 5-12). Lane 13, the DNA marker.

### A unique RNA structure feature on pre-mRNAs is associated with polyadenylation and alternative polyadenylation

Another key step of mRNA processing is polyadenylation that starts with endonucleolytic cleavage on pre-mRNAs followed by addition of a poly(A) tail at the cleavage site^2^. Since only the pre-mRNA structure before endonucleolytic cleavage can be used for elucidating the mechanism underpinning polyadenylation, we assessed whether pre-mRNAs before endonucleolytic cleavage were enriched in our nuclear SHAPE-Structure-Seq libraries. We compared the sequencing reads coverage across cleavage sites (poly(A) sites) annotated in a previous study^34^ with both our nuclear SHAPE-Structure-Seq libraries and cytosolic SHAPE-Structure-Seq libraries. The reads across poly(A) sites were highly enriched in our nuclear SHAPE-Structure-Seq libraries compared to our cytosolic SHAPE-Structure-Seq data (Supplementary Fig. 9). This indicated high enrichment of pre-mRNAs before polyadenylation in our nuclear SHAPE-Structure-Seq libraries (Supplementary Fig. 9).

To accurately determine RNA structure features across poly(A) sites, only reads mapped across poly(A) sites and in downstream flanking regions were used to generate SHAPE reactivity profiles (3,077 and 551 poly(A) sites with >=1 RT-stop per nucleotide under *in vivo* and *deproteinized* conditions were used in the analysis, Supplementary Data 6 and 7). We found that average SHAPE reactivities in two regions (from −28 nt to −17 nt upstream of the poly(A) site and from −4 nt to +1 nt across the poly(A) site) were significantly higher compared to flanking regions for both *in vivo* and *deproteinized* conditions (Fig. 5a,b), suggesting these two regions tended to be more single-stranded than flanking regions. To eliminate the effect of nucleotide composition, we identified control sites where nucleotide composition was similar to the sequence content across poly(A) sites, but where polyadenylation did not occur (Supplementary Fig. 10a,b). We found no significant RNA structure features across these control sites, indicating the two single-stranded regions observed across the poly(A) sites above, were specifically associated with polyadenylation (Fig. 5a,b). Furthermore, we assessed whether these two single-stranded regions also appeared in alternative polyadenylation sites. Compared to constitutive poly(A) sites, we found a similar but weaker structure feature across alternative polyadenylation sites (Supplementary Fig. 11a,b, Supplementary Data 8 and 9). Notably, these structure features were different to those identified from a previous RNA structurome study on mature mRNAs^24^, further indicating structure differences between pre-mRNAs and mature mRNAs. Therefore, this RNA structure feature with two single-stranded regions may also be responsible for alternative polyadenylation.

**Fig. 5.**
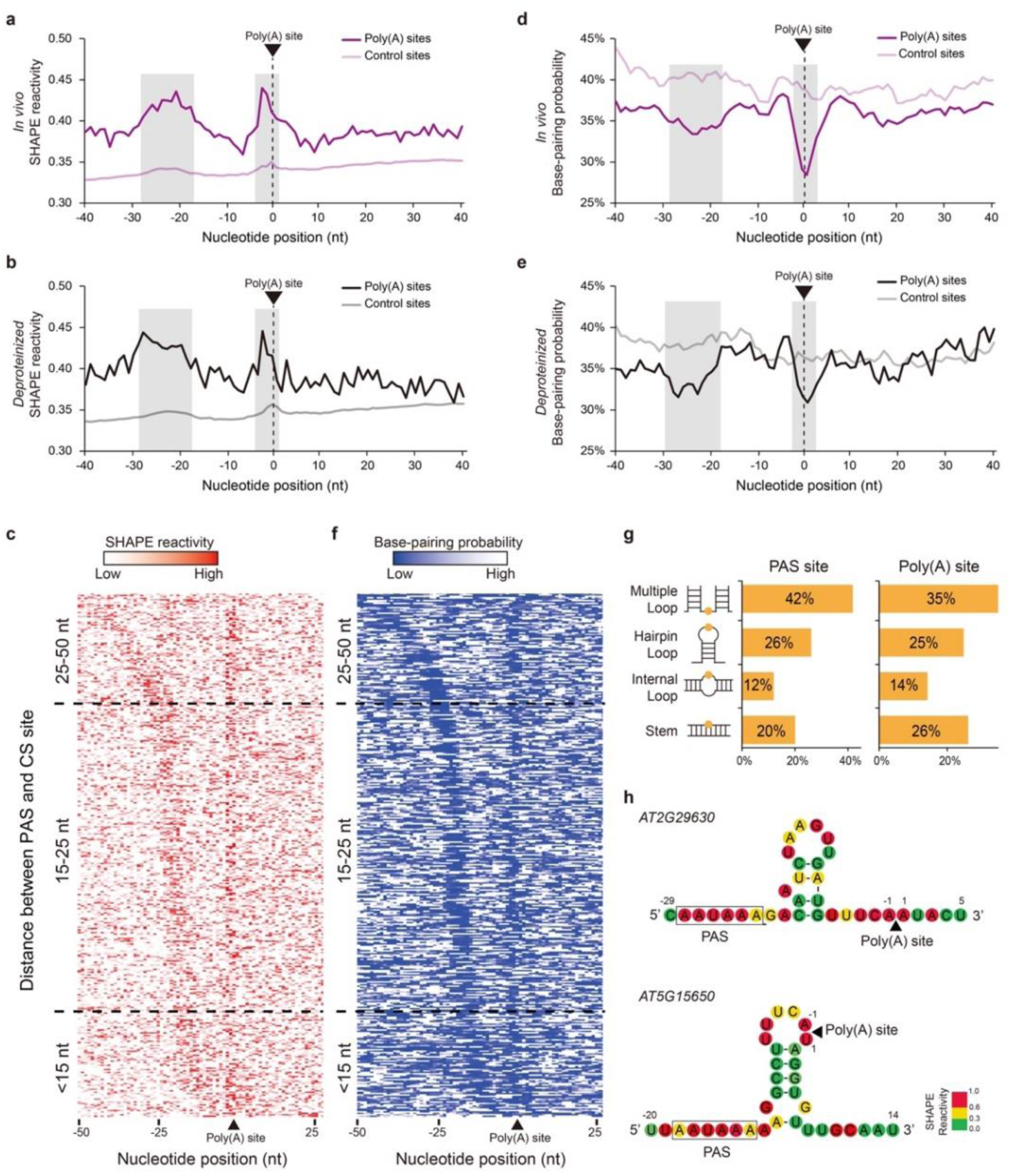
Two single-stranded regions on pre-mRNA are associated with polyadenylation. **a,b**, SHAPE reactivity profiles across poly(A) sites for *in vivo* (**a**) and *deproteinized* (**b**) conditions. The X-axis represents the relative position to the poly(A) site. Average SHAPE reactivities in two regions (from −28 nt to −17 nt upstream of the poly(A) site and from −4 nt to +1 nt position across the poly(A) site) were significantly higher compared to flanking regions for both *in vivo* (purple) and *deproteinized* (black) conditions (Fisher’s exact test, *P*-value = 3.1e-14 and 3.6e-6 for *in vivo; P*-values = 4.7e-12 and 1.4e-3 for *deproteinized*). The corresponding average SHAPE reactivity profiles for the control sites are in light colours. **c**, Heatmap showing *in vivo* SHAPE reactivity profiles across the PAS motif “AAUAAA” and poly(A) site. The pre-mRNAs are sorted by the distance between PAS and poly(A) site. The gradient colour from light to dark red represents SHAPE reactivity from low to high. The SHAPE reactivities are much higher at both the PAS and poly(A) sites compared to flanking regions. **d, e**, Base-pairing probability (BPP) profiles across poly(A) sites for *in vivo* (**d**) and *deproteinized* (**e**) conditions. Average BPPs in two regions (from −28 nt to −17 nt upstream of the poly(A) site and from −4 nt to +1 nt position across the poly(A) site) were significantly lower compared to flanking regions for both *in vivo* (purple) and *deproteinized* (black) conditions (Fisher’s exact test, *P*-value = 5.0e-12 and 1.4e-7 for *in vivo; P*-values = 1.3e-8 and 2.8e-6 for *deproteinized*). The corresponding average BPPs for the control sites are in light colours. **f**, Heatmap showing *in vivo* BPPs across the conventional PAS motif “AAUAAA” and poly(A) site. **g**, Classification of RNA structure elements across the PAS and poly(A) sites. The three different single-stranded types (multiple loop, hairpin loop and internal loop) and the double-stranded stem type were assessed for all the PAS and poly(A) sites. The percentage of each type is shown. Most of the PAS and poly(A) sites are located in the single-stranded loop regions including multiple loop, hairpin loop and internal loop. **h**, Illustrations of two individual pre-mRNA structures with both the PAS and poly(A) sites located in single-stranded loop regions. Nucleotides were colour-coded according to the *in vivo* SHAPE reactivity values (SHAPE reactivity 0.6-1.0 marked in red, 0.3-0.6 marked in yellow, 0-0.3 marked in green).

Further investigation of the sequence content in positions −28 nt to −17 nt upstream of poly(A) sites showed that this region had an accumulation of the conventional polyadenylation signal (PAS) motif “AAUAAA” (Supplementary Fig. 12, Supplementary Data 10). We then aligned SHAPE reactivities across this conventional PAS motif “AAUAAA” upstream of poly(A) sites and sorted pre-mRNAs by the distance between PAS and poly(A) sites (Fig. 5c). The corresponding SHAPE reactivities across PAS and poly(A) sites for each pre-mRNA were then plotted as a heatmap (Fig. 5c). We found that SHAPE reactivities were higher at both PAS sites and across poly(A) sites compared to flanking regions (Fig. 5c). Thus, the conventional polyadenylation signal (PAS) motif “AAUAAA” tended to be a single-stranded region. Interestingly, this unique structure feature consistently appeared regardless of the distance between PAS and poly(A) sites (Fig. 5c). Hence, our results suggested that the single-strandedness of both PAS and poly(A) sites may serve as RNA structure signals for polyadenylation.

To understand what type of RNA structures could be formed with these two single-stranded regions, we folded sequences across the poly(A) sites with the constraints of SHAPE reactivities by using the *Vienna RNAfold* package^35^. We then calculated the base-pairing probability (BPP) of each nucleotide^35^. Consistent with our SHAPE reactivity profiles, we found that the BPPs in these two regions (from −28 nt to −17 nt upstream of the poly(A) site and from −4 nt to +1 nt across the poly(A) site) were significantly lower compared to the flanking regions for both *in vivo* and *deproteinized* conditions, confirming the single-strandedness of these two regions (Fig. 5d,e). Furthermore, we found no obvious BPP features across the control sites, indicating this structure feature was not due to preferential nucleotide composition (Fig. 5d,e). We also generated the heatmap of BPPs across the conventional PAS motif “AAUAAA” and poly(A) sites. We found that the BPPs were much lower at both PAS sites and poly(A) sites compared to flanking regions (Fig. 5f), consistent with SHAPE reactivity profiles (Fig. 5c). In addition, we assessed the detailed RNA structure elements across PAS and poly(A) sites using the *Forgi* utility^36^. We found that most RNA structures had both PAS and poly(A) sites located in single-stranded loop regions including multiple loop, hairpin loop and internal loop (Fig. 5g). For instance, one type of RNA structure comprised both PAS and poly(A) sites located in multiple loop regions and connected by one hairpin structure (an example is illustrated in Fig. 5h-top). Another type of RNA structure comprised the PAS site located in a multiple loop region with the poly(A) site located in a hairpin loop region (an example is illustrated in Fig. 5h-bottom). Therefore, our results indicated that diverse RNA structures were formed to maintain single-strandedness at both PAS and poly(A) sites.

## Discussion

For the first time, we generated the *in vivo* RNA structure landscape of *Arabidopsis* nuclear RNAs with structure information for all four nucleotides by developing nuclear SHAPE-Structure-Seq. Having achieved high coverage and high accuracy with our nuclear SHAPE-Structure-Seq, we were able to investigate global RNA structure features of nuclear mRNAs and uncover the regulatory role of RNA structure in mRNA processing.

### Nuclear mRNA structures are globally different from cytosolic mRNA structures

Cytosolic mRNAs are the processed products from nuclear mRNAs, thus they share the same sequences. An intriguing question is whether nuclear mRNA structures in these regions are the same as cytosolic mRNA structures? A recent study in mammalian cells showed that nuclear mRNAs were generally more folded than cytosolic mRNAs^21^. We found similar phenomena in our study (Fig. 2a). However, by further dissecting different genic regions, we found that nuclear mRNA structures in exons located in UTR regions tended to be more single-stranded than cytosolic mRNA structures (Fig. 2b). Previous individual RNA structure studies showed that strong RNA structures in 5’UTRs of mature mRNAs are required for recruitment of translation initiation factors^37^. Also, strong RNA structures in 3’UTRs are critical for mature mRNA stability^38^. Structure differences between nuclear and cytosolic mRNAs at 5’ UTRs and 3’ UTRs might be associated with translation initiation and mature mRNA stability. Furthermore, we observed that nuclear mRNAs tended to be more folded in CDS regions compared to cytosolic mRNAs (Fig. 2b). Since ribosomes are known to remodel mature mRNAs by unwinding RNA structures^19,39^, more single-stranded features in cytosolic mRNA coding regions may be caused by ribosome scanning. In other words, nuclear mRNA structures without interference from ribosomes may remain more folded.

From our global to local assessment of RNA structure features, we found that RNA structure features downstream of start codons and at stop codons were significantly different between nuclear and cytosolic mRNAs. A previous *in vitro* study suggested that mature mRNAs might require strong structures downstream of the start codon for increasing the 40S subunit “dwell time”^37^. Our observation (Fig. 2c) implied that stronger structures downstream of the start codon in cytosolic mRNAs compared to nuclear mRNAs might relate to the ribosome pausing *in vivo*. At stop codons, we found much higher SHAPE reactivities in cytosolic mRNAs (Fig. 2d). This single-stranded structure feature was also observed in a previous RNA structurome study and was suggested to facilitate translation termination^40^. But in nuclear mRNAs, this structure feature was much weaker (Fig. 2d), implying this single-stranded structure feature at stop codons in cytosolic mRNAs might be specific for translation termination. Taken together, these structure feature differences between nuclear and cytosolic mRNAs implied that mRNAs might undergo refolding from the nucleus to the cytosol.

In addition to the effects on structure differences from translation, mRNA processing, e.g. polyadenylation and splicing, might also impact the folding status of RNA structures in different cellular compartments. Previous RNA structure profiling of mature mRNAs after polyadenylation in human observed more folded structure features in the region downstream of PAS sites compared to the region upstream of PAS, which were found to facilitate polyadenylation^15^. However, we did not observe significant structure differences between these two regions in our nuclear SHAPE-Structure-Seq, suggesting mRNAs might be refolded after polyadenylation (Fig. 5a,b). In addition, we found a distinctive single-stranded region across poly(A) sites (Fig. 5a,b), demonstrating that our method had overcome the limitations of previous mature mRNA structurome studies, which lacked structure information across poly(A) sites^15^. Furthermore, our previous study on mature mRNAs in *Arabidopsis* revealed that significantly more folded structure features formed upstream of alternative polyadenylation sites compared to flanking regions^24^. However, we found RNA structure features associated with alternative polyadenylation in the pre-mRNAs before polyadenylation (Supplementary Fig. 10a,b) were different from those observed in mature mRNAs^24^. Additionally, our previous study on mature RNAs showed a stronger RNA structure feature upstream of 5’ss in unspliced events ^24^. However, we did not observe similar features in our nuclear SHAPE-Structure-Seq (Fig. 3a,b), indicating the RNA structure features related to splicing are also different between pre-mRNAs and mature mRNAs^24^. Thus, these structure differences before and after mRNA processing implied that mRNAs may adopt different structures for serving distinct biological processes. Many other factors, e.g. diverse protein interactions, RNA modifications and distinct cellular conditions between the nucleus and cytosol, may also contribute to these structure differences, which offers scope for future studies.

### Distinctive RNA structure features upstream of 5’ss and at the branch point are associated with recognizing 5’ss and 3’ss respectively

Distinct from mammalian splicing where exon skipping is the dominant type of alternative splicing, intron retention is the most common alternative splicing event in plants and can result in significant biological consequences^41^. Previous *in vitro* enzymatic RNA structure profiling in *Arabidopsis* nuclear RNAs showed greater structure differences at the exon-intron junctions where the 5’ end of introns were much more double-stranded than upstream exons and 3’ end of introns were more single-stranded than flanking sequences^14^. However, we did not observe these dramatic differences across exons and introns in our nuclear SHAPE-Structure-Seq data, further confirming that *in vivo* RNA structures were different from *in vitro* RNA structures^19,20^.

The recognition of both 5’ss and 3’ss are of great importance during splicing^1,4^. The consensus sequence motifs for both are so short that a large number of sites with matching sequences are widely spread in the transcriptome^4^. How to distinguish actual splice sites from a large number of false positives has been a primary challenge in elucidating the regulation of splicing^4^. Previous individual studies in human suggested strong RNA structures at U1 and U2 snRNA binding sites can prevent the interactions with U1 and U2 snRNA, thus interfering with the recruitment of U1 and U2 snRNPs during splicing^42–44^. In our transcriptome-wide study for 5’ss, we identified a two-nucleotide single-stranded RNA structure feature immediately upstream of the 5’ss, which was associated with splicing events (Fig. 3a,b). Since the structure feature was located within the U1 snRNA binding region (from −3 to +6 position across the 5’ss)^33^, it is likely that the single-strandedness of these two nucleotides promotes the binding of U1 snRNA in 5’ss recognition. For 3’ss, we found the single-strandedness at the branch point was associated with splicing events (Fig. 3a,b). Since U2 snRNA binds across the branch point through base-pairing^1^, the single-strandedness at the branch point might promote the binding of U2 snRNA in 3’ss recognition. Alternatively, this single-strandedness might also be a consequence after binding with U2 snRNA since the RNA-RNA base-pairing interaction leaves the branch point as an internal bulge^1^. Previous studies in yeast suggested that stem-loop structures between the branch point and 3’ss could promote the recognition of 3’ss^45,46^. We also found a 4nt low SHAPE reactivity region upstream of AG dinucleotides at the 3’ss, which suggested the formation of a stronger RNA structure between 3’ss and the branch site (Fig. 3a,b). However, this structure feature was not associated with splicing events, and as such, might be linked with subsequent steps after the recognition of 3’ss, such as docking the 3’ss into the reaction center to approach 5’ss^47^. Notably, the two-nucleotide single-stranded RNA structure feature upstream of 5’ss and the single-strandedness at the branch point were also strongly associated with the selection of alternative 5’ss and 3’ss, respectively (Fig. 3c,d,e,f). These results further suggested that these two *in vivo* RNA structure features might serve as general rules for determining actual 5’ss and 3’ss in splicing. Although we observed global SHAPE reactivity difference between *in vivo* and *deproteinized* libraries (Supplementary Fig. 5), we found very similar structure features across splicing sites under these two conditions (Fig. 3). A previous biophysics study suggested that splicing occurred rapidly once splice sites were recognized^48^. Therefore, our result suggested that the *in vivo* RNA structure features we observed across splicing sites might represent the RNA structures of pre-mRNAs before spliceosome assembly.

### The two-nucleotide single-stranded RNA structure feature upstream of 5’ss can regulate splicing

Previous studies of individual RNA structure suggested that strong RNA structures formed at 5’ss can inhibit U1 snRNA binding, and subsequently repress splicing^8,43,44^. However, the strong structures in each case were so different that no general RNA structure features have been identified for regulating splicing. From our nuclear SHAPE-Structure-Seq data, we were able to sensitively determine that a very fine RNA structure feature showing single-strandedness at the −1 and −2 positions upstream of 5’ss was associated with splicing at the transcriptome-wide scale (Fig. 3a,b). Our functional assessment further confirmed that fine-tuning RNA structure by switching the base-pairing status of only these −1 and −2 positions upstream of 5’ss was sufficient to change the fate of splicing (Fig. 4).

One possible mechanism is the single-strandedness of the −1 and −2 positions upstream of 5’ss promoted splicing by facilitating the binding of U1 snRNA. U1 snRNA base-pairs with a total of nine nucleotides (from −3 to +6 region of 5’ss) across 5’ss^33^. Thus, any nucleotides within this nine-nucleotide U1 binding site should have been able to affect splicing. However, we observed that single-strandedness at all other nucleotide positions within the U1 binding site (except for the −1 and −2 positions) were not able to rescue splicing events (Fig. 4b,c). Therefore, our study revealed that the position of this two-nucleotide single-stranded RNA structure feature was also important for regulating splicing. This phenomenon raised the possibility that the −1 and −2 nucleotides upstream of 5’ss may be the first positions for the interaction with U1 snRNA. Further biophysics studies might be able to assess this hypothesis. Furthermore, once the 5’ss is recognized by base-pairing with U1 snRNA, the whole spliceosome is assembled onto the intron region and the 5’ss-U1 interaction is replaced by interactions of 5’ss with U5 (from −3 to −1 region of 5’ss) and U6 (from +4 to +8 region of 5’ss) snRNAs^49^. It is possible that the single-strandedness of the −1 and −2 positions may also promote interaction with U5 snRNA. Taken together, both our transcriptome-wide RNA structure profiling and functional assessment indicated that the two-nucleotide single-stranded structure feature at the −1 and −2 positions upstream of 5’ss can serve as a general role in splicing regulation.

Since splicing is a fundamental biological process across eukaryotes, the regulatory motif for splicing is likely to be conserved and highly selected during evolution. Previous identification of the most conserved sequence motif required for 5’ss recognition is as short as only a dinucleotide GU at 5’ss^4^. The sequence requirement of only two nucleotides might be minimized during evolution selection. The short sequence length of the conserved nucleotides might provide the plasticity for flanking nucleotides to contribute to other biological functions. Here, we postulate that the very fine RNA structure feature we identified from the transcriptome is likely to have evolved in a similar way as the sequence motif, in terms of the single-strandedness of only two nucleotides being sufficient to regulate splicing. It will be of great interest to extend our study in other species to investigate the generality of this regulatory mechanism.

### Two single-stranded regions upstream and across poly(A) sites are associated with both polyadenylation and alternative polyadenylation

Similar to the challenge of how to recognize splice sites, the recognition of poly(A) sites does not always rely on sequence content. In particular, no unique sequence motif exists around poly(A) sites in plants^11,50^. Indeed, only ∼10% of *Arabidopsis* genes contain the conventional PAS motif “AAUAAA” upstream of poly(A) sites^11^. Therefore, how to precisely determine actual poly(A) sites has been a major question for improving our understanding of polyadenylation regulation. A previous enzymatic probing study on *in vitro* nuclear RNAs in *Arabidopsis* had attempted to investigate RNA structure features at poly(A) sites^14^. However, no structure features were observed at either polyadenylation or alternative polyadenylation sites^14^, which may be due to the low resolution of enzymatic probing or low comparability of single-stranded and double-stranded RNase probing^6,14^. Here, we identified two single-stranded regions (from −28 nt to −17 nt upstream of the poly(A) sites and from −4 nt to +1 nt across the poly(A) sites) that were associated with both polyadenylation and alternative polyadenylation (Fig. 5a,b,d,e and Supplementary Fig. 11). These RNA structure features did not appear in the regions where the nucleotide composition was similar but polyadenylation did not occur (Fig. 5a,b,d,e). Hence, these close-by two single-stranded RNA structure features may serve as an additional signature for the recognition of poly(A) sites.

Interestingly, most conventional PAS motifs “AAUAAA” are located within the region from −28 nt to −17 nt upstream of the poly(A) sites (Supplementary Fig. 12). We did observe the conventional PAS motif “AAUAAA” region was more single-stranded compared to flanking regions (Fig. 5c,f), which suggested that the single-stranded region upstream of the poly(A) site corresponded to the PAS motif site. Since sequence content is insufficient for predicting PAS sites^11^, the single-stranded region upstream of poly(A) sites could offer another signature for recognizing the unconventional PAS motif. Moreover, the interactions of the PAS sites with CPSF30 and WDR33 proteins are crucial during polyadenylation^2^. Hence, PAS sites might adopt this single-stranded structure feature to facilitate protein binding. Furthermore, the endonucleolytic cleavage at poly(A) sites is catalyzed by CPSF73, which has been suggested to prefer RNA single-strandedness^51^. Therefore, the single-stranded region across poly(A) sites might facilitate the interaction between CPSF73 and poly(A) sites.

In summary, we generated the *in vivo* nuclear RNA structure landscape in *Arabidopsis* achieving both high resolution and accuracy with our nuclear SHAPE-Structure-Seq method. We revealed both global and local structure differences between nuclear and cytosolic mRNAs. We successfully identified respective pre-mRNA structure features associated with splicing and polyadenylation. Through functional validation we determined an RNA structure feature which can regulate splicing. Our study unveiled a new RNA structure regulatory mechanism for mRNA processing. Also, our work emphasized the importance of dissecting RNA populations from different stages of the mRNA life cycle in order to investigate the relationship between RNA structure and biological functions.

## Acknowledgements

This research was supported in part by the NBIP Computing infrastructure for Science (CiS) group through the provision of a High-Performance Computing Cluster. We thank Prof. Peter Shaw and Prof. Chun Kit Kwok for their advice on the experimental design. We are also grateful to Prof. Igor Vorechovsky for discussions. This work was supported by the Biotechnology and Biological Sciences Research Council [BB/L025000/1], the Norwich Research Park Science Links Seed Fund and a European Commission Horizon 2020 European Research Council (ERC) Starting Grant [680324].

## Author contributions

Y.D. conceived the research and designed the experiments; Q.L., X.Y., Y.Z., and X.C. performed the experiments; Z.L. designed the data analysis and experimental validation; Z.L., M.N., and J.C. performed the data analysis with assistance from Y.D.; Z.L. and Y.D. wrote the manuscript with input from all authors. Z.L. and Q.L. contributed equally to this work.

